# Energy expenditure does not explain step length-width choices during walking

**DOI:** 10.1101/2021.04.11.439375

**Authors:** Stephen A. Antos, Konrad P. Kording, Keith E. Gordon

## Abstract

Healthy young adults have a most preferred walking speed, step length, and step width that are close to energetically optimal. However, people can choose to walk with a multitude of different step lengths and widths, which can vary in both energy expenditure and preference. Here we further investigate step length-width preferences and their relationship to energy expenditure. In line with a growing body of research, we hypothesized that people’s preferred stepping patterns would not be fully explained by metabolic energy expenditure. To test this hypothesis we used a two-alternative forced-choice paradigm. Fifteen participants walked on an oversized treadmill. Each trial participants experienced two stepping patterns and then chose the pattern they preferred. Over time, we adapted the choices such that there was 50% chance of choosing one pattern over another (equally preferred). If people’s preferences are based solely on metabolic energy expenditure, then these equally preferred stepping patterns should have equal energy expenditure. We found that energy expenditure differed across equally preferred step length-width patterns (p < 0.001). On average, longer steps with higher energy expenditures were preferred over shorter and wider steps with lower energy expenditures (p < 0.001). We also asked participants to rank a set of shorter, wider, and longer steps from most preferred to least preferred, and from most energy expended to least energy expended. Only 7/15 participants had the same rankings for their preferences and perceived energy expenditure. Our results suggest that energy expenditure is not the only factor influencing a person’s conscious gait choices.

## Background

Most healthy individuals do not lunge, shuffle, or waddle when they walk. There are some gait patterns that we prefer more than others. A person’s most preferred gait speed, step length, and step width are close to energetically optimal ^1–4^. Just as each gait pattern has an associated energetic cost, each gait pattern also has a perceived preference. Similar to how studying how people control their gait, understanding how people perceive their gait may help identify what factors drive movement decisions.

Utility theory provides a framework to examine people’s preferences and decision making ^5,6^. A utility function can measure preferences over a set of movement parameters. Such functions are often represented as cost functions or loss functions (negative of utility functions) for optimization problems ^7,8^. Classic examples of cost functions for upper extremity reaching tasks include minimizing torque ^9^, jerk ^10^, or variation in endpoint error ^7^. Utility functions can also be applied to conscious decision via a two-alternative forced-choice paradigm. That is, a person experiences two movement choices and then selects the movement they preferred. For example, you could be given a choice to run or walk 400 meters. If you cared most about minimizing cost of transport, you should choose to walk^11^. Whereas if you cared minimizing the amount of time it takes, you should choose to run. Applying utility theory to walking can help us describe the choices one makes, and further understand what factors influence a person’s movement decisions.

We can examine a person’s step length-width choices and test whether their choices are energy optimal because people can voluntarily walk with different, non-preferred step lengths and widths. Based on previous studies, a person’s most preferred step length and width coincides with an energetic minimum ^2,4,12^. However, this evidence only considers a single point in their utility function: the point of maximum utility (Figure 1A). Because energy expenditure can often explain walking behaviors, we chose to investigate whether energy expenditure was equal for equally preferable stepping patterns. Walking with shorter, wider, or longer than normal steps can affect both energy expenditure ^4,12^ and a person’s preference. At non-preferred step lengths and widths, other factors (e.g. stability^13,14^, joint torques, muscle length^15^) influencing a person’s decisions may also become amplified and shift their preferences away from the energetic optimum.

**Figure 1-.**
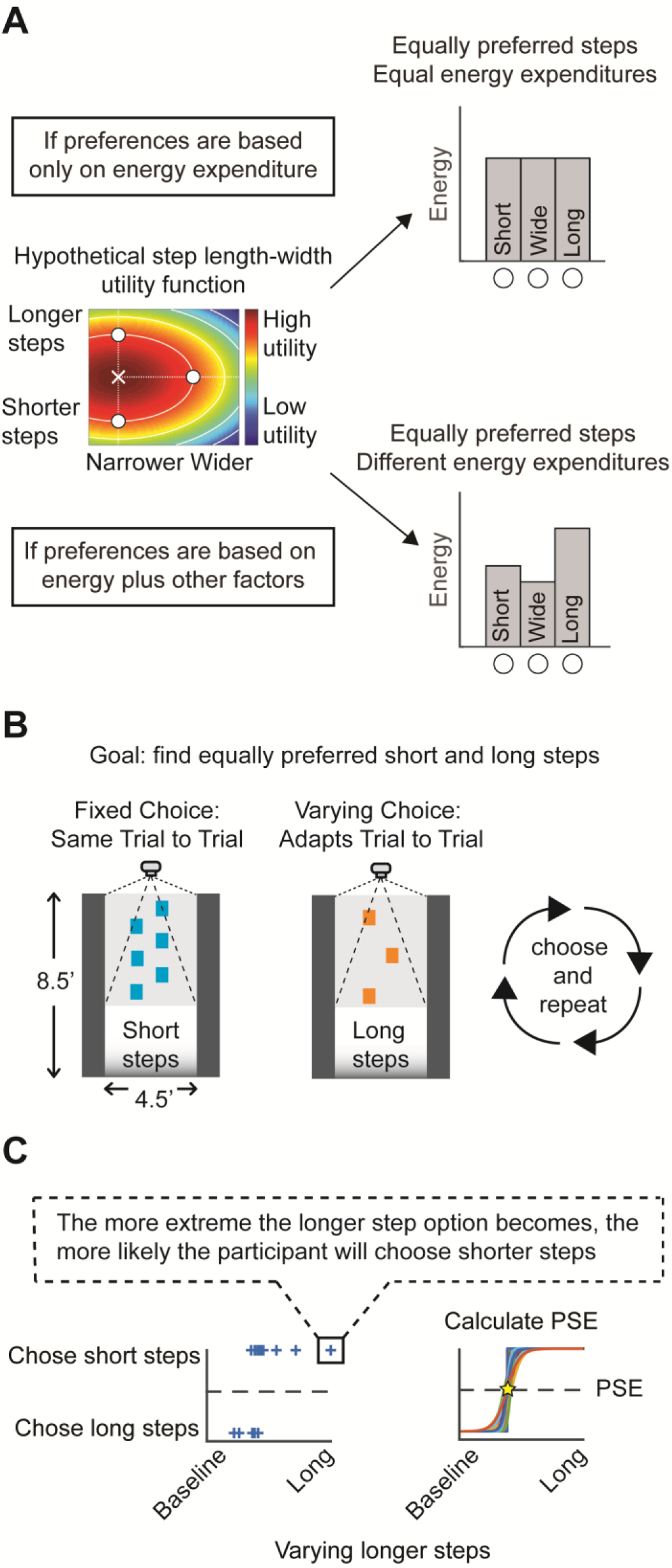
Experimental methods. A) Hypothetical step length-width utility function. The “X” is a person’s most preferred step length-width pattern, and has the highest perceived utility. The white contours are indifference curves – curves of equal utility. The white circles are indifference points and lie on one indifference curve. If a person’s preferences are based solely on energy expenditure, then energy should be equal along indifference curves. B) Example to find a pair of equally preferred shorter and longer steps. Participants walked with two stepping patterns: shorter than normal steps (fixed every trial), and longer than normal steps (varied each trial). The participant chose the stepping pattern they preferred most, and walked with that pattern again. The longer step option adapts on future trials to make the decision more difficult on subsequent trials. C) Example of choice data: a blue cross represents a participant’s decision. For example, when longer steps became more extreme, the participant preferred shorter steps, and vice versa. We bootstrapped the decision data, fit logistic curves, and calculated the point of subjective equivalence (PSE), where *p*=0.5.

Here we used a two-alternative forced choice paradigm to find sets of step length-width patterns that were perceived as equally preferable, and then measured energy expenditure for each pattern. If energy expenditure dominates a person’s decisions, then energy expenditure should be equal across equally preferred step length-width patterns. If energy expenditure is not equal across equally preferred step length-width patterns, there must be other factors dominating a person’s choice of step length and width (Figure 1A). We hypothesized that energy expenditure would be different across equally preferable step length-width patterns.

## Methods

### Participants

We recruited fifteen healthy young adults between the ages of 18-30; participants were excluded if they had any injuries or impairments that affected their gait or decision making. This study was approved by the Northwestern Institutional Review Board (STU00203331) and all participants provided written, informed consent.

### Experimental design

Our experiment consisted of two parts: 1) finding a set of equally preferred step length-width patterns and 2) measuring energy expenditure for the equally preferred set. Participants walked on an oversized treadmill (Tuff Tread, Willis, TX) at a constant speed (1.2 m/s), while a projector (Hitachi, Japan, 60Hz) displayed rectangles as stepping targets to control step length and step width. A black poster board was placed in front of the treadmill so participants could plan at least two steps ahead. We collected kinematic data using 11 active markers and a 12 camera motion capture system (Qualisys, Sweden). We placed markers on the calcaneus, second metatarsal, fifth metatarsal, lateral malleolus, greater trochanter, and T10 spinous process. Markers were labelled in real time and streamed at a rate of 60Hz to a custom Matlab program (Mathworks, Natick, MA, version 2013B) to provide feedback to the experimenter about participants’ step length and width. Step length and width were calculated as the maximum distance between fifth metatarsal markers for each step. In the second part of the experiment, we measured the 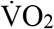 using indirect calorimetry (K4b^2^, Chicago, IL) to calculate metabolic power.

### Finding equally preferred step length-width patterns

Participants began by walking on the treadmill for at least 5 minutes to acclimate to our experimental setup, and then for two additional minutes as we measured their baseline (most preferred) step length and width. Next, participants practiced walking on rectangular targets for five minutes to become comfortable walking with different step lengths and widths. We used a two-alternative forced choice paradigm to find a set of equally preferred shorter, wider, and longer than normal steps (Figure 1B). Equally preferred patterns, also known as indifference patterns, are step length-width patterns in which there is a 50% chance of choosing one pattern over the other. Each trial, participants walked with two stepping patterns for 10 seconds each. Then participants chose the pattern they preferred most and walked with that pattern again for 10 seconds. We fixed one of the two choices to be 75% of the participant’s baseline step length (short steps) and 100% of their baseline step width. The other choice in each trial was either a wide or long step option that adapted based on the participants’ previous decisions. For the wide step option, we fixed step length to the participant’s baseline step length while we varied the step width. For the long step option, we fixed step width to the participant’s baseline step width while we varied the step length. We chose short steps to be the fixed pattern because, in our pilot studies, all participants were capable of taking much shorter steps as opposed to much longer steps. Participants completed 30 trials for each preference pair (short-wide and short-long) for a total of 60 trials. For each preference pair, we bootstrapped the participants’ decisions (N=10,000), fit logistic curves, and found the point of subjective equivalence (p = 0.5, Figure 1C).

We provided participants with the following instructions: “After you have experienced both choices, please tell us which one you preferred the most (choice one or choice two) and you will walk with it again. If you cannot physically perform one of the choices, you can walk normally during that choice and should choose the other option. We are studying walking, so one foot should always be in contact with the ground. Please do not jump.”

We pre-randomized the order of the wide and long step options over all trials, as well as the choice order (one or two) within each trial. We wanted participants to step accurately, but also wanted the task to feel natural. After each trial, we calculated the step by step error by subtracting the actual step length and width from the desired step length and width. If participants’ average step length or width error over all steps for that trial was greater than 3cm, we provided the participant with a verbal cue to make their steps wider, narrower, shorter, or longer depending on the direction of the error.

### Measuring energy expenditure

After finding equally preferred shorter, wider, and longer than baseline steps, we measured 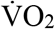 (mL/s), normalized by body weight (kg), and calculated metabolic power (W/kg) for six conditions. We first collected trials for standing, baseline walking with no stepping targets, and then baseline walking with stepping targets. Participants then walked with the equally preferred shorter, wider, and longer steps in a random order. All trials were six minutes: the first three minutes were for the participant to reach steady state and the last three minutes were used to calculate the average 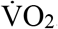. We then calculated metabolic power using the Brockway Equation ^16^. We subtracted the metabolic power of the standing trial from all walking trials to obtain the net metabolic power (W). Respiratory exchange ratios 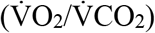 indicated predominately aerobic metabolism for all subjects and conditions.

### Subjective Reports

We were also interested in participants’ subjective decision-making process, whether there was a change in stepping preferences across time scales (10 seconds vs. 6 minutes), and how well participants could perceive their energy expenditure. After the finding equally preferred shorter, wider, and longer steps, participants commented on their decision-making process. After walking with each pattern for six minutes during metabolic data collection, we asked participants to rank order the shorter, wider, and longer steps from most preferred to least preferred, and from most energetically costly to least energetically costly.

### Statistical Analysis

We used a linear mixed effects model to test whether metabolic power was different for equally preferred step length-width patterns. Our dependent variable was metabolic power. We treated the type of stepping pattern (shorter, wider, and longer) as a fixed effect; participants were treated as random effects (intercept only).

We also performed a descriptive analysis for participants’ subjective preference rankings, subjective energy rankings, and actual energy expenditure. First, for each participant, we calculated whether the percent difference in energy expenditure for equally shorter, wider, and longer steps. We used a percent difference tolerance of 10% to determine whether energy expenditures were “approximately equal”. This tolerance is greater than the reported effects of arm swing during running^17^ Then, we examined whether participants’ subjective preference and subjective energy rankings matched. Finally, we looked at whether the subjective preference and subjective energy rankings aligned with actual energy expenditure. Rankings were considered “not aligned” only if there was a meaningful difference for that participant. That is, if a participant had similar energy expenditures (<10% difference), their rankings would always be considered “aligned” with actual energy expenditure.

## Results

Fifteen participants (7 Female /8 Male, age 25.3 + 2.5 years) walked on an oversized treadmill while stepping onto projected visual targets to guide their step length and width. Each trial, participants walked with two step length-width patterns, chose the pattern they preferred, and walked with that pattern again. We found a set of equally preferable shorter, wider, and longer than normal steps and then measured energy expenditure for each step length-width pattern. If a person’s decisions were based solely on energy expenditure, then energy expenditure should be equal for equally preferred stepping patterns.

### Step length-width preferences

Participants could perform the stepping task with an average error less than 3cm, and we could find a set of equally preferred shorter, wider, and longer than normal steps for all participants. We found that stepping preferences varied across participants (Figure 2).

**Figure 2 –.**
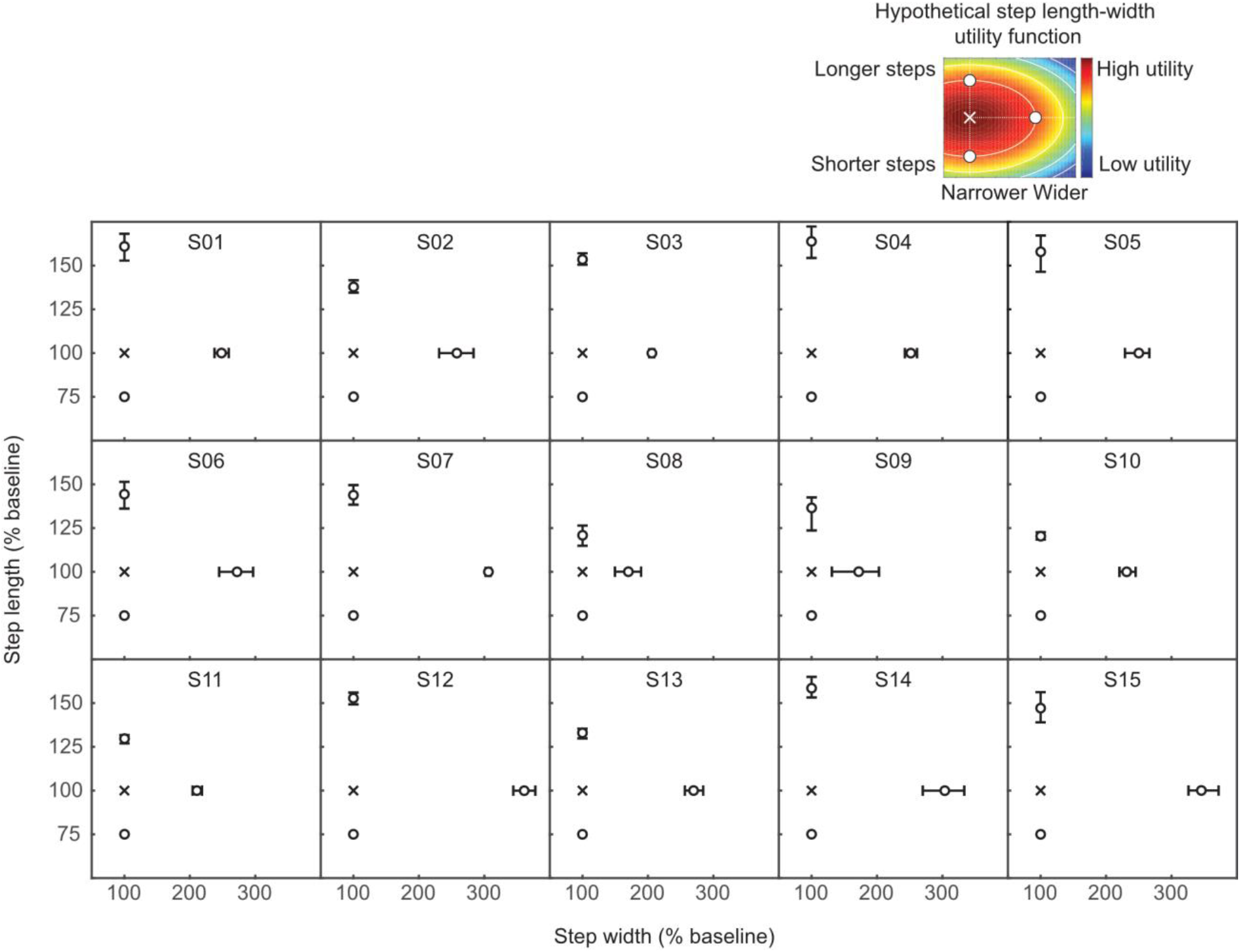
Equally preferred step length-width patterns. The X denotes a participant’s baseline step length and step width; circles denote a set of equally preferred shorter, wider, and longer than baseline stepping patterns. Error bars denote 95%confidence intervals.

### Energy expenditure for equally preferred step length-width patterns

To determine whether energy expenditure was the only factor influencing step length-width preferences, we tested if equally preferred shorter, wider, and longer than normal steps had equal energy expenditures. We found that metabolic power was not equal for equally preferred step length-width patterns (Figures 3 and 4, p < 0.001, F_2,42_ = 16.2). Longer steps had greater metabolic power than the shorter (p < 0.001, F_1,42_ = 30.8) and wider steps (p < 0.001, F_1,42_ = 15.1); there was no significant difference between shorter and wider steps (p = 0.10, F_1, 42_ = 2.8). Only one participant had energy expenditures that were similar across all patterns (<10% difference). Energy expenditure could not fully explain participants’ step length-width decisions.

**Figure 3 –.**
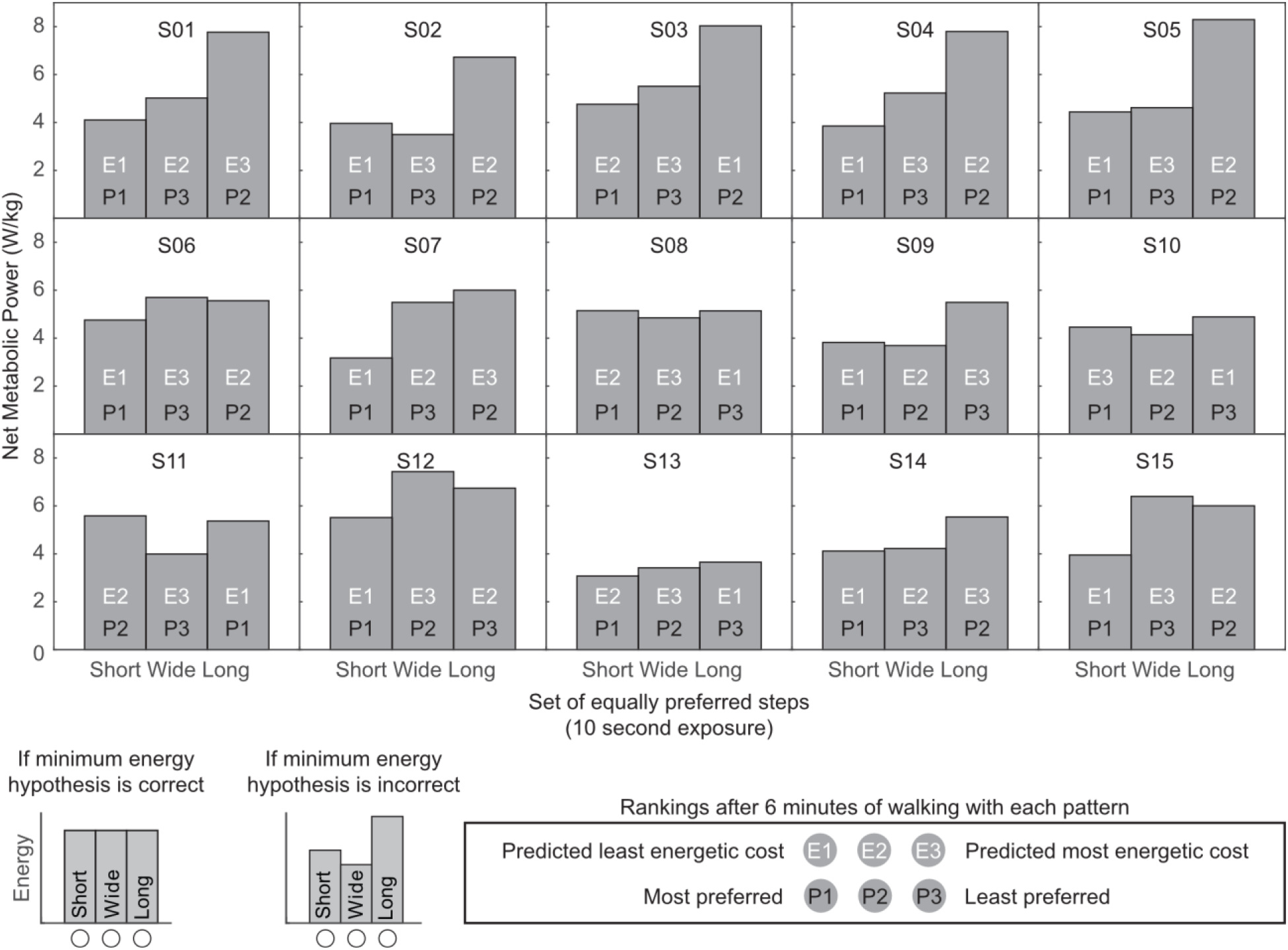
Net metabolic power for equally preferred step length-width patterns for 10 second trials. Only one participant (S08) had net metabolic powers that had less than a 10% difference across their equally preferred short, wide and long steps. Participants also ranked the set of equally preferred stepping patterns after a longer exposure time (six minutes) from 1) most preferred to least preferred and 2) least energetic cost to most energetic cost. Subjective preference and energy rankings after six minutes of walking did not match for 8/15 participants. Furthermore, of the participants with matching preference and energy rankings (7/15), only three had rankings that aligned with their actual metabolic power.

**Figure 4 –.**
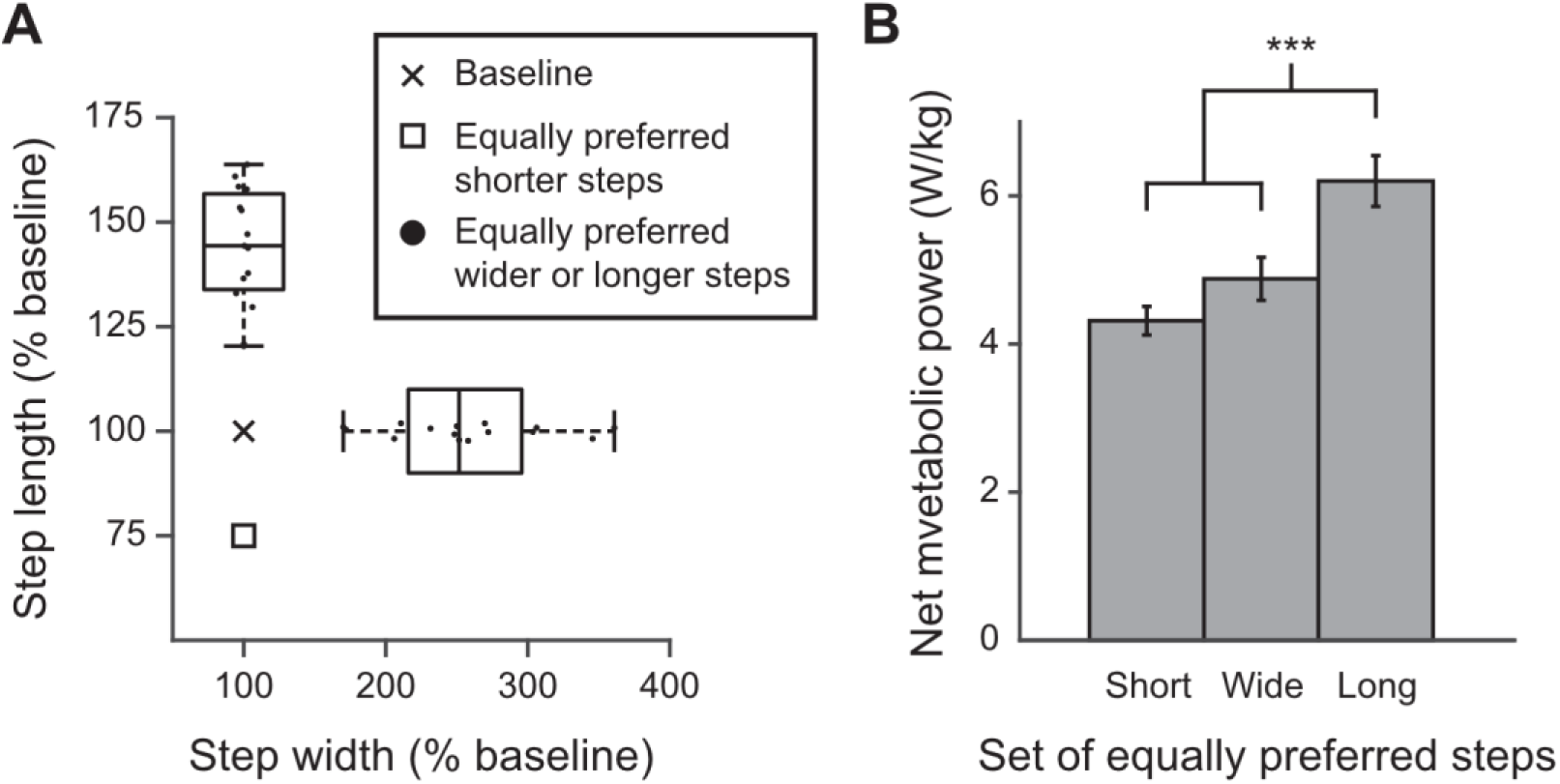
Grouped results. A) Box and whisker plots for equally preferred shorter, wider, and longer than normal steps across all participants. There was a large range of preferences across participants. B) Average net metabolic power for equally preferred shorter, wider, and longer than normal steps over all participants. Error bars denote 95% confidence intervals. On average, participants preferred longer steps at an increased energy expenditure over shorter and wider steps (p < 0.001).

### Subjective rankings

After collecting metabolic data, we asked participants to rank the shorter, wider, and longer than normal steps from most preferred to least preferred, and from most energetically costly to least energetically costly. Of the 15 participants, only 7 had subjective energy and preference rankings that matched, 8 had preference rankings that aligned with actual energy expenditure, and 8 had energy rankings that aligned with actual energy expenditure (Table 1). All but one participant preferred shorter steps the most when walking for six minutes. Two thirds of participants preferred longer steps over wider steps, often reporting that the wider steps were less comfortable or required increased hip effort. Even on a longer time scale, subjective rankings and objective measurements of energy expenditure did not align with preferences for many participants.

**Table 1 –.** Preference and energy rankings after six minutes with each pattern

| Participant | Were energy expenditures approximately equal? | Did preference/energy rankings match? | Did preference rankings align with energy expenditure? | Did energy rankings align with energy expenditure? |
| --- | --- | --- | --- | --- |
| 1 | N | N | N | Y |
| 2 | N | Y | N | N |
| 3 | N | N | N | N |
| 4 | N | Y | N | N |
| 5 | N | Y | N | N |
| 6 | N | Y | Y | Y |
| 7 | N | N | Y | Y |
| 8 | Y | N | Y | Y |
| 9 | N | Y | Y | Y |
| 10 | N | N | Y | N |
| 11 | N | Y | N | N |
| 12 | N | N | Y | Y |
| 13 | N | N | Y | N |
| 14 | N | N | N | Y |
| 15 | N | Y | Y | Y |
| **Total Y** | 1/15 | 7/15 | 8/15 | 8/15 |

## Discussion

In this study we investigated if a person’s step length-width preferences could be explained by energy expenditure. First, we found a set of equally preferred shorter, wider, and longer than normal steps for healthy young adults. Then, we tested whether energy expenditure was equal for equally preferred stepping patterns. We found that step length-width preferences did not align with energy expenditure, suggesting that additional factors influence a person’s stepping preferences. We also found that preferences changed between short (10 seconds) and long (6 minutes) time scales. For all but one participant, shorter steps became more preferable for longer bouts of walking. Although preferences changed for longer bouts of walking, many participants’ preferences still did not align with their actual or perceived energy expenditure.

We demonstrated how to find equally preferred step length-width patterns using a two-alternative forced choice paradigm. Our approach can be extended to other movement parameters that people can voluntarily control (e.g. gait speed) to investigate people’s choices as a function of movement. Few studies have explicitly measured movement preferences ^18–21^, particularly for walking ^11^. Ultimately, the goal of many rehabilitation interventions is to change a person’s movement behavior. If we can understand which factors most influence a person’s movement preferences, then targeting those factors may, hopefully, lead to greater changes in behavior. Based on the variability in preferences and personal anecdotes from our participants, we believe that these factors are likely person specific.

We found that energy expenditure alone cannot explain a person’s step length-width preferences. While several studies have found that people naturally select the most energetically efficient gait^1,3,22–24^, there is also growing body of evidence demonstrating that people do not always adopt the most energetically efficient gait^11,13,25–27^. For example, when walking downhill, people naturally select a gait pattern that increases stability but also increases metabolic cost.^13^ Our study differs from the studies above because we examined participants’ movement perception rather than movement control. In our study, participants made conscious choices based on their perceived movement preference, rather than being asked to perform a task and naturally adopting gait pattern. This distinction is critical because perception (i.e. preference) and control (action) utilize different neural mechanisms, and by measuring movement perceptions we cannot make conclusions about movement control. Studying movement perception, control, and how they interact is critical to further our understanding of walking behavior.

There were several limitations to our study. While we have shown that energy alone cannot explain preferences, we cannot make conclusions about what other factors influence decisions, or the extent their influence. We limited our decision trials to 10 seconds because it provided participants enough time to make consistent decisions but was short enough to make our experiment feasible. Although participants’ decisions remained consistent during preference measurements, preferences are likely dynamic and can change over different time scales. Furthermore, we had participants walk with novel gaits. People may need more experience or longer exposure times to be able to make metabolically efficient decisions for unfamiliar gait patterns. However, we found that many participants could not correctly rank energy expenditure after six minutes of exposure, and their stepping preferences after six minutes often differed from the actual and perceived energy expenditures.

Previous work suggests that time ^11^, comfort ^25^, and stability^13^ may also contribute to a person’s gait choices. We held time constant across all choices in our study, so while time and energy may both influence a person’s choice of gait, even these factors combined cannot fully explain step length-width preferences. Several of our participants reported that wide steps were their least preferred pattern – even though they were more energetically efficient – because it was uncomfortable for their hip abductors. This suggests that the perception of local factors may contribute more to movement utility than global energy expenditure. This was unsurprising, given that many of our participants could not correctly rank energy expenditure for the set of equally preferred short, wide, and long steps.

### Conclusion

We demonstrated how to measure equally preferred stepping patterns and then found that equally preferable gaits do not translate into energy minimization.

## Acknowledgements

The authors would like to thank Geoffrey Brown for his discussions and feedback related to this study.

